# Polymorphic 30-nm Chromatin Fiber and Linking Number Paradox. Evidence for the Occurrence of Two Distinct Topoisomers

**DOI:** 10.1101/478396

**Authors:** Davood Norouzi, Victor B. Zhurkin

**Affiliations:** Laboratory of Cell Biology, CCR, National Cancer Institute, NIH, Bethesda, MD 20892, USA

## Abstract

We discuss various models of spatial organization of the 30-nm fiber that remains enigmatic despite 40 years of intensive studies. Using computer simulations, we found two topologically different families of the fiber conformations distinguished by the linker length, L: the fibers with L = {10*n*} and {10*n*+5} bp have DNA linking numbers per nucleosome *DLk »* –1.5 and –1.0, respectively. The fibers with *DLk »* –1.5 were observed earlier, while the topoisomer with *DLk »* –1.0 is novel. These predictions were confirmed for circular nucleosome arrays with precisely positioned nucleosomes. We suggest that topological polymorphism of chromatin fibers may play a role in the process of transcription, which is known to generate different levels of DNA supercoiling upstream and downstream from RNA polymerase. In particular, the {10*n*+5} DNA linkers are likely to produce transcriptionally competent chromatin structures. This hypothesis is consistent with available data for several eukaryotes, from yeast to human. We also analyzed two recent studies of chromatin fibers – on the nucleosome crosslinking *in vitro,* and on radioprobing DNA folding in human cells. In both cases, we show that the novel topoisomer with *DLk »* –1.0 has to be taken into account to interpret experimental data. This is yet another evidence for occurrence of two distinct fiber topoisomers. Potentially, our findings may reflect a general tendency of chromosomal domains with different levels of transcription to retain topologically different higher-order structures.

## Introduction

Eukaryotic chromosomes are hierarchically organized into distinct substructures with specialized functions. Recent chromosome conformation capture [1-4] and super-resolution microscopy studies [5,6] have demonstrated partitioning of chromosomes are into multi-scale loops, from the left-handed nucleosome supercoils, up to the megabase topological domains. At each level of the hierarchy, the chromatin folding patterns are thought to be related to various genomic processes such as transcription and replication.

In the past decade, enormous amount of valuable information has been revealed about 3D organization of DNA at the both ends of this hierarchical chain. At the local end, several X-ray [7] and Cryo-EM [8] structures of oligonucleosomal arrays were resolved, with and without linker histone. At the mega scale end, large DNA loops were observed, stabilized by nucleoprotein complexes involving CTCF and cohesin (in interphase chromosomes) or condensin (in metaphase) [9-11]. However, the intermediate level (corresponding to ~100 nm) remains mostly in the shadow.

Notably, the 3D organization of the so-called 30-nm fiber is still under debate [12-13], even in the relatively simple case of regular nucleosome arrays studied *in vitro*. In particular, it is questionable whether the one-start (solenoid) fiber with bent DNA is formed for the long inter-nucleosome linkers, L ≥ 50-60 bp [14-16] or various types of two-start and multi-start folds with relatively straight linker DNA are predominant for all nucleosome repeat lengths (NRLs) [17-19]. More importantly, it remains unclear if and how this 3D folding is related to regulation of transcription *in vivo* [5, 20].

Below, we briefly describe our recent findings related to DNA topology in the 30-nm fiber, and present new evidence for the existence of two distinct topoisomers of chromatin fibers characterized by different nucleosome spacing.

## Topological polymorphism of chromatin fibers

Soon after the nucleosome was discovered [21-23], it became obvious that there is a discrepancy between the nucleosome structure and DNA topology in SV40 minichromosome (the so-called “DNA Linking Number Paradox” first formulated by Crick [24] and Fuller [25]). This conundrum has been described many times during the past 40 years [26,27], so we can skip technical details. In short, measurements of DNA topology in circular minichromosomes in viruses showed generation of only one superhelical turn per nucleosome [28,29], instead of 1.6-1.7 negative turns expected from the left-handed DNA wrapping in nucleosome.

(The DNA linking number, Lk, defines the number of times each DNA strand winds around the other. For closed circular DNA, the change in the linking number, ΔLk (compared to the relaxed state of DNA), the change in DNA Twisting, ΔTw, and DNA Writhing, Wr, are related by the well-known equation: ΔLk = ΔTw + Wr.)

To explain the above paradox, several models of the 30-nm fiber, with *D*Lk = –1 and –2, have been suggested in the 1980-ies [30-32]. They remained untested for about 20 years, until the first crystal structure of tetranucleosome has been solved by Richmond and coworkers [7]. This structure, as well as the Cryo-EM and X-ray fiber conformations published later [8], were obtained for the tandem repeats of strongly positioned nucleosomes ‘601’ [33].

However, the linking number paradox still remained unresolved, mainly for two reasons. First, because in all these fiber structures, the DNA trajectories are similar, with *D*Lk estimated to vary between –1.4 to –1.3 [34], which differs significantly from *D*Lk = –1.0 measured for SV40 minichromosome [28,29] and later, for the plasmids containing 5S DNA repeats [35]. Second, in the published X-ray and cryo-EM structures of oligonucleosomal arrays [7,8] the linkers contain integral number of DNA turns (L=20, 30, 40 bp, denoted here as {10n}), whereas biochemical [36,37] and genomic studies [38-40] show prevalence of the linkers with half-integral numbers of DNA turns (denoted as {10n+5}). Therefore, it was critically important to clarify whether the chromatin fibers with {10n} and {10n+5} linkers fold differently, as this might be a key to resolve the ΔLk paradox.

To address this issue, we systematically analyzed stereochemically feasible two-start fibers, with the linker lengths varying in a wide interval from 10 to 70 bp [18,34]. As a result, we discovered, *in silico*, two distinct families of fiber conformations (T2 and T1) characterized by different DNA topologies. One family, T2, is represented by topoisomers similar to the fibers observed experimentally [7,8], while the other family, T1, contains novel forms with a different DNA folding (Figures 1A, B). Importantly, the topology of a chromatin fiber strongly depends on the nucleosome spacing: the energetically optimal fibers with the {10n} and {10n+5} linkers have the DNA linking numbers per nucleosome *D*Lk *»* –1.5 and –1.0, respectively. It should be noted that the predicted change in DNA linking number, *DD*Lk *»* 0.5, is actually a strong difference – it means that shifting two nucleosomes in a circular array by 5 bp changes the number of DNA supercoils by one.

**Figure 1.**
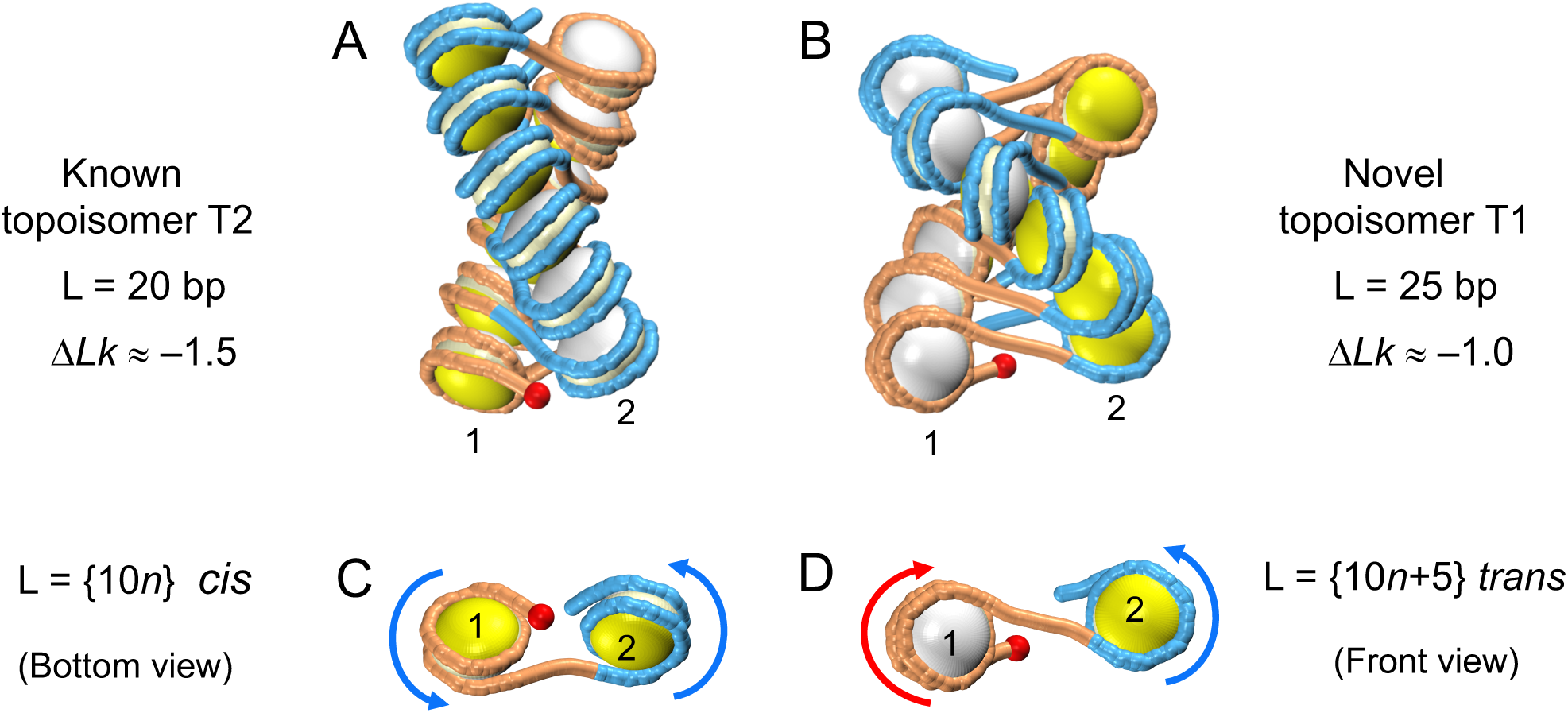
Distinct ‘rotational setting’ of adjacent nucleosomes in the T2 and T1 topoisomers produces different conformations of the nucleosome arrays. **(A, B)** Energetically optimal two-start fibers with L=20 and L=25 bp, respectively [18]. Note that the red spheres (indicating the entry points in the ‘first’ nucleosome) are positioned differently in the two fibers: in (A) they are facing the viewer, while in (B) they are located in the rear half of the nucleosome #1. In each nucleosome, the entry side is colored in yellow, and the exit side is in white. **(C, D)** The nucleosomes #1 and #2 of the fibers in (A, B) are shown. **(C)**Bottom view. Both nucleosomes are facing the viewer with the yellow side, and the arrows indicating the DNA trajectory are rotated similarly, counter-clockwise. L = {10n}. The DNA linker contains (approximately) integral number of helical turns of DNA, thus, the two adjacent nucleosomes are in a cis-like configuration. **(D)**Front view. Note the different orientations of the nucleosomes #1 and #2: they are facing the viewer with the white and yellow sides, respectively. The arrows are rotated clockwise (#1) and counter-clockwise (#2). L = {10*n*+5}. Changing the linker DNA length by 5 bp introduces additional half-turn of the DNA duplex, resulting in a trans-like configuration of the two nucleosomes.

We obtained this result rigorously [18,34], evaluating the Wr and *D*Lk values through calculations of the Gauss double integral [41,42]. At the qualitative level, it is illustrated in Fig. 1, where the difference in *D*Lk between the T2 and T1 topoisomers is reduced to alteration of the ‘rotational setting’ of adjacent nucleosomes by 180° which is a consequence of changing the DNA linker length by 5 bp, from {10n} to {10n+5}. The change in the linker DNA twisting by 180° corresponds to *DD*Lk *»* 0.5 mentioned above, therefore, intuitively we can link the topological difference between the T2 and T1 forms to the cis-to trans-like transition shown in Figures 1C, D.

The predicted topological polymorphism of chromatin fibers was confirmed experimentally [19], by employing topological gel assay and EM imaging. Using circular DNA constructs with regularly positioned nucleosomes ‘601’ we demonstrated that the nucleosome arrays with NRL = 167 and 172 bp are characterized by *D*Lk = –1.4 and –0.9, respectively [19]. In other words, the DNA supercoiling changes by as much as 50% depending on the length of the DNA linker between nucleosomes, in excellent agreement with theoretical results [18]. This observation was made for relatively short 20- and 25-bp linkers observed in yeast [39,40]. Recently, we corroborated this conclusion analyzing nucleosome arrays with 183- and 188-bp NRL [43], typical for higher eukaryotes (L = 36 and 41 bp).

Thus, we made an important step toward resolving the long-standing Linking Number Paradox. We have proven that there is no single *D*Lk value characterizing ensembles of various chromatin fiber configurations <u>*in general*</u>. In fact, the average linking number is defined by nucleosome spacing and varies at least from –0.9 to –1.4. (According to computations [18,34] *D*Lk varies from –0.8 to –1.7 in the energetically feasible conformations.) The value *D*Lk = –1.26 measured for the yeast minichromosomes by Roca *et al*. [27] fits in this interval and reflects the average *D*Lk for those configurations of the relatively short nucleosome chains that were stabilized under the experimental conditions.

In addition to affecting the topology of a nucleosome array, the length of linker DNA also mediates the compactness of chromatin folding. Sedimentation analysis of the nucleosome ‘601’ arrays with variable linker lengths allowed Grigoryev and coworkers [17] to observe a periodic modulation of chromatin folding with strongest changes between the compact arrays with L ={10n} and unfolded nucleosome chains with L = {10n+5}. These experimental findings are generally consistent with our initial computations [18] indicating that the nucleosome stacking interactions in the arrays with {10n} linkers are more favorable compared to {10n+5} linkers, and thus, the former arrays are likely to fold in a more stable and compact configuration than the latter. A more detailed comparison was made recently. Our Monte Carlo simulations incorporating dynamic unwrapping of nucleosomal DNA [44] proved to be in a very close quantitative agreement with sedimentation velocity measurements [17] for a wide range of arrays (L = 20 – 60 bp) effectively covering NRLs in most eukaryotic genomes.

In summary, our data suggest that both topology (or level of DNA supercoiling) and compactness of chromatin fibers are defined by the linker length (and therefore, by NRL).

## Nucleosome spacing and the level of transcription

The topological polymorphism of chromatin fibers described above may play a role in regulation of transcription. According to the model of Liu and Wang [45] the level of negative supercoiling of DNA is decreased downstream and increased upstream of the transcription complex. Therefore, we hypothesized that existence of the two types of fibers (T1 and T2) with different linking numbers may be related to the transient DNA topological changes occurring during transcription. We reckoned that the T1 topoisomer with *D*Lk *»* –1 (and a weak supercoiling of DNA) would be formed predominantly downstream from RNA polymerase (in the highly transcribed genes), as opposed to the T2 topoisomer with *D*Lk *»* –1.5, which is likely to be stabilized in the upstream regions (and more generally, in the regions with a low level of transcription).

Since the T1 and T2 topoisomers are characterized by distinct linker lengths, {10n+5} and {10n}, respectively, we expected to see a difference in the distribution of sizes of inter-nucleosome linkers in the highly and lowly expressed genes. To verify this assumption, we compared the nucleosome positions in the yeast genes from the top and bottom 25% of the expression level scale [34]. Indeed, the two sets of genes were found to have different distributions of nucleosome repeats: for the highly expressed genes NRL ≈ 161 bp (the average linker length <L> = 14 bp), and for the lowly expressed genes NRL ≈ 167 bp (<L> = 20 bp).

These results are consistent with the above hypothesis that nucleosomal arrays with L ≈ 10*n*+5, which have a relatively low superhelical density, are transcriptionally more competent than the arrays with L ≈ 10*n*. In addition, the fibers with L ≈ 10*n*+5 reveal a greater plasticity [17], which may facilitate formation of gene loops [46] and enhancer-promoter loops [47], thereby further inducing transcription of the corresponding genes. By contrast, in the inactive genes the prevalent linker length is L ≈ 10*n* which corresponds to a higher superhelical density and a higher stability of the chromatin fiber.

In higher eukaryotes, the hierarchical genomic organization is more complicated than in yeast, and it is unlikely that there are simple ‘mechanistic’ ways of modulating gene expression by changing the 3D folding of chromatin at the level of nucleosome arrays. Nevertheless, available data suggest that there are certain correlations between the DNA linker lengths and chromatin epigenetic states [48,49], which are essential for gene regulation, among other genomic processes. In particular, the H3K9-methylated constitutive heterochromatin regions in Drosophila have the average linker length <L> = 30 bp, or {10n}, while the polycomb-repressed H3K27-methylated chromatin has <L> = 26 bp [49]. Importantly, the highly expressed genes display the shortest linkers, with average <L> = 17 bp – that is, close to {10n+5} values. In mouse embryonic stem cells [50], the genome-wide mapping of nucleosomes revealed the preferential {10n+5} linkers, L=35 and 45 bp (on average, it gives <L> = 40 bp, which is consistent with NRL = 196 bp reported earlier [51]). By analogy with yeast [34] and fly [49], for ~20% of the most actively transcribed mouse genes the distribution of the linker lengths is shifted toward the smaller L values, so that on average <L> = 35 bp (D.N and V.B.Z., unpublished observation).

Above, we presented available data supporting a hypothetical relationship between DNA topology, inter-nucleosome spacing and the level of expression of corresponding genes. Next, we proceed to two recent studies of chromatin fibers. The first one is on the nucleosome crosslinking *in vitro* [52], and the second one is on radiation-induced cleavage of DNA *in situ*, in human cells [53]. In both cases, experimental data cannot be interpreted based on the published X-ray and Cryo-EM structures of chromatin fibers [7,8], and therefore, some alternative fiber conformations have to be taken into account. We show that the novel T1 topoisomer described above is likely to be such an alternative.

## The linker length dependence of nucleosome–nucleosome crosslinks *in vitro*

In their landmark 2004 study, Richmond and coworkers [54] used Cys-Cys disulfide bonds to stabilize and visualize the two-start chromatin fiber with NRL = 177 bp. Recently, Ekundayo *et al.* [52] further extended the disulfide crosslinking approach to analyze specific nucleosome– nucleosome contacts in chromatin fibers in solution. In addition to the mutants H2A-E64C and H4-V21C used by Dorigo *et al.* [54] they prepared several H2B mutants, including H2B-Q44C (Figure 2A). In this way, very interesting results were obtained, suggesting that the histone–histone contacts (and, therefore, the chromatin fiber configuration) vary significantly with NRL. In particular, a clear distinction was observed between the H4-H4 and H2B-H2B crosslinking patterns for the fibers with {10n} and {10n+5} linkers. The H4-H4 crosslink was notably stronger for {10n} linkers (NRL = 157, 167 and 177 bp), while the H2B-H2B crosslink was detected only for {10n+5} linkers (NRL = 162 and 172 bp), see Figure 2B.

**Figure 2:**
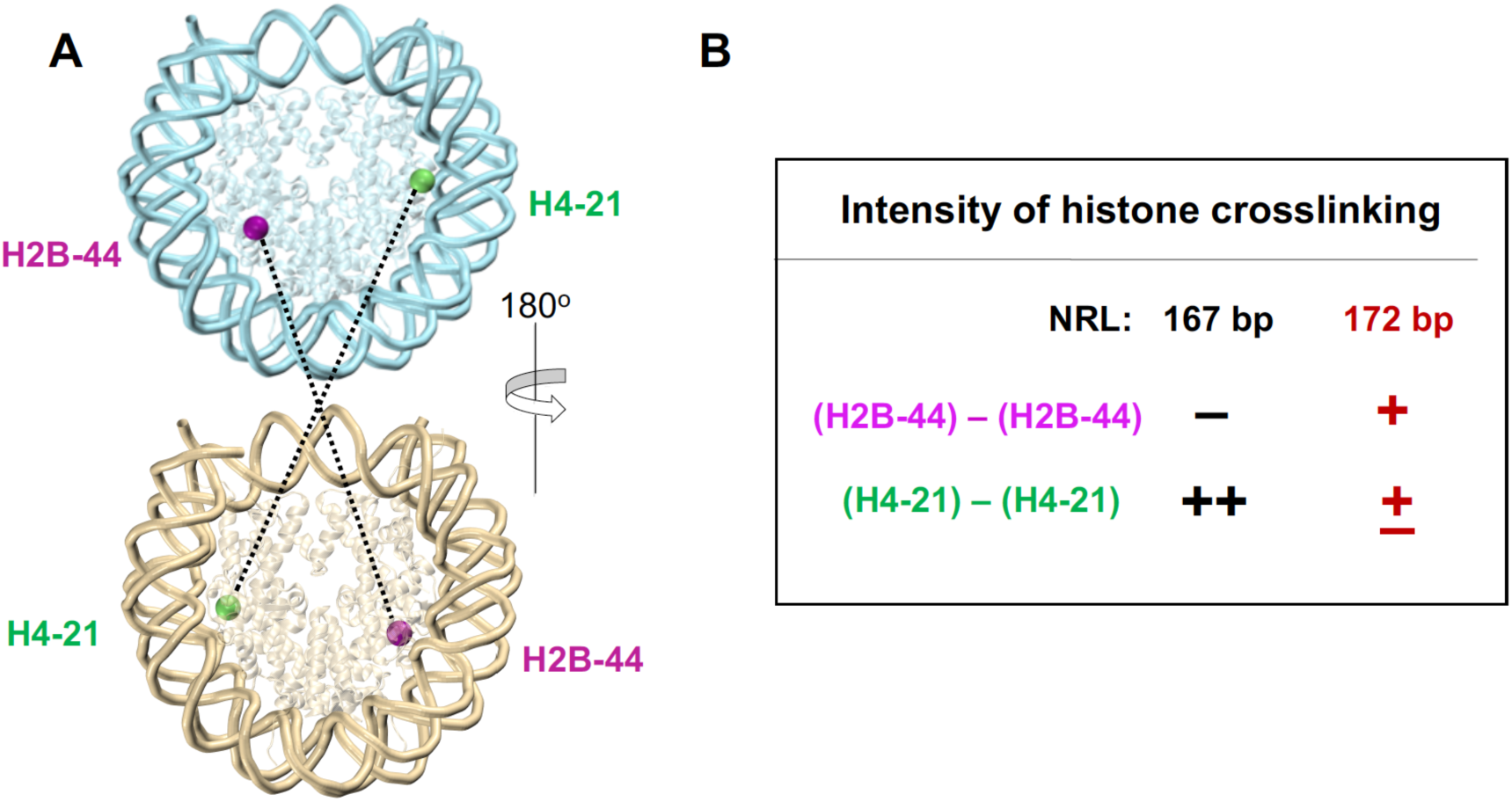
Specific nucleosome–nucleosome contacts in chromatin fibers in solution analyzed with disulfide crosslinking [52]. **(A)** The ventral and dorsal views of nucleosome core particle showing the Cys residues chosen for crosslinking in sphere representation (H2B-Q44C: magenta, H4-V21C: green). **(B)** Intensity of crosslinks strongly depends on the nucleosome repeat. The nucleosome arrays containing 12 repeats of strongly positioned sequence ‘601’ [33] were used. The crosslinked signals measured by Ekundayo *et al.* [52] are shown in a qualitative representation. Results for NRL = 162 bp, and NRL = 157, 177 bp are similar to the presented data for NRL = 172 and 167 bp, respectively (Figures 5 and S4 in [52]).

This difference cannot be explained based on the available X-ray and cryo-EM structures [7,8], because all of them belong to the {10n} type. Therefore, we analyzed the stereochemically possible fiber structures that were calculated earlier [18,34]. First, we realized that the energetically optimal structures (with strong nucleosome stacking) are not suitable for formation of either the H4-H4 or the H2B-H2B crosslinks (Figure 3, center). Next, we found that crosslinking between the two H2B-44 residues requires lateral displacement of the “top” nucleosome to the left, whereas the H4-H4 crosslink is only possible when the “top” nucleosome is shifted to the right (Figure 3). The lateral displacements strongly correlate with the nucleosome inclination angle *r*, the parameter used to distinguish between the two-start fiber topoisomers T1 and T2 [18].

**Figure 3:**
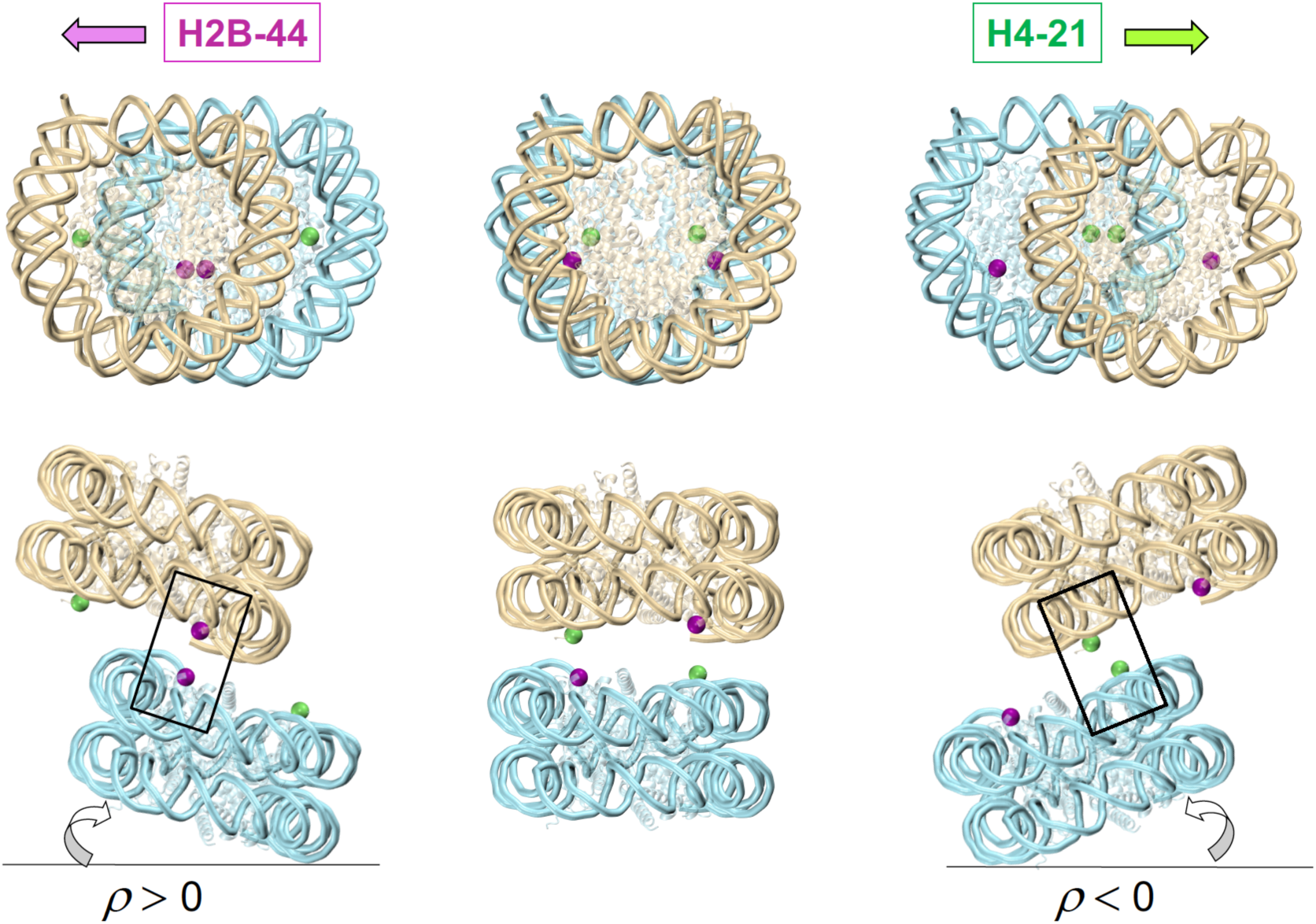
Formation of disulfide crosslinks requires spatial rearrangement of nucleosomes. The face-to-face interaction between two nucleosomes is shown (the top view above and the side view below). Central panel: The strong overlap between nucleosomes (that is favorable energetically) is not consistent with the disulfide crosslinking H4-H4 or H2B-H2B. Left panel: Crosslinking between two H2B-44 residues (magenta) requires lateral displacement of the “top” nucleosome to the left which is accompanied by positive inclination of nucleosomes, *r* > 0. By contrast, crosslinking between two H4-21 residues (green) requires lateral displacement of the “top” nucleosome to the right which correlates with negative inclination, *r* < 0 (right panel).

These conformations are energetically unfavorable, which agrees with the experimental observation that the intensities of H4-H4 and H2B-H2B crosslinks are relatively low, compared to the H4-H2A crosslinks used by Dorigo *et al.* [54] to stabilize the ‘canonical’ two-start fiber with NRL = 177 bp. Therefore, to analyze such displaced conformations we performed Monte Carlo (MC) simulations [44], instead of energy optimization used earlier to describe the energetically optimal structures [18].

By generating the MC ensemble of conformations, we obtained distribution of the distances between the H2B-44 residues in nucleosomes (i) and (i+2) facing each other in the two-start fibers (Figure 3, left panel). As follows from Figure 4A, the probability of the two H2B-44 residues being close to each other is significantly higher for the fibers with NRL = 172 bp. In this case, the distance H2B-H2B is less than 30 Å in 10% of fiber conformations in the MC ensemble, while for NRL = 167 bp, only 1% of MC conformations satisfy this criterion. The latter result is in a qualitative agreement with Figure 2B, indicating that the H2B-H2B crosslinks were not detected for NRL = 167 bp.

**Figure 4.**
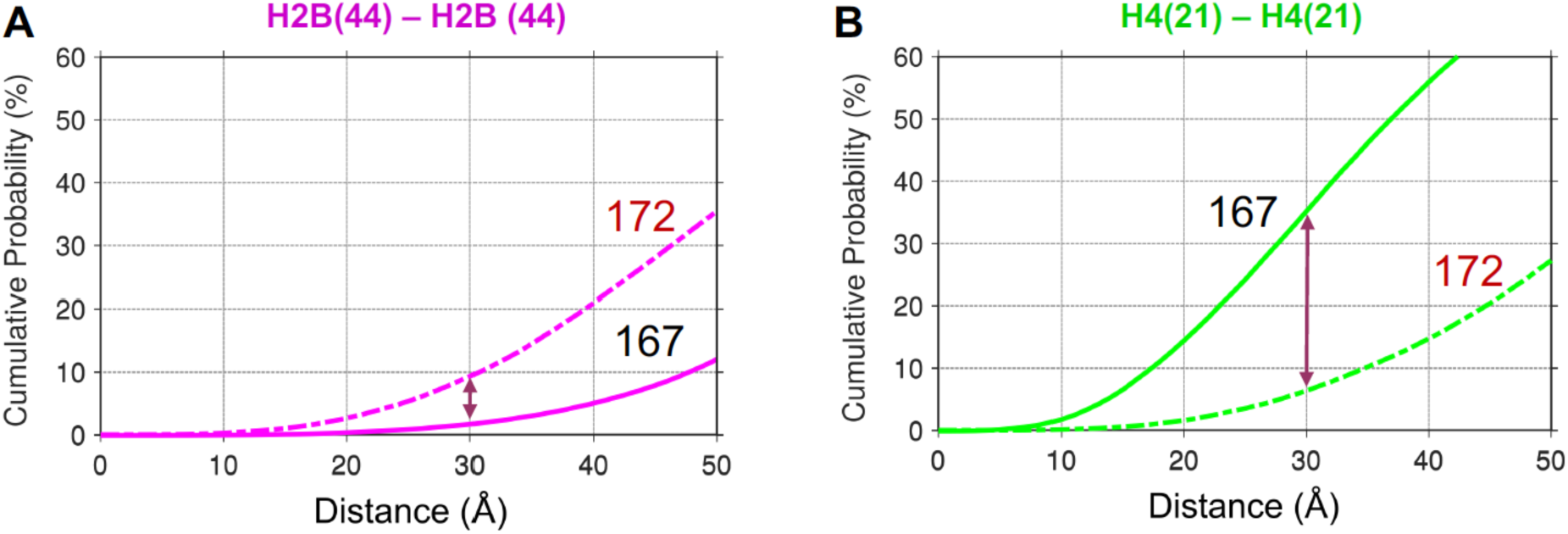
Distribution of the H2B-H2B **(A)** and H4-H4 **(B)** distances between the nucleosomes (i) and (i+2) in the two-start fibers. Monte Carlo ensembles of 100,000 conformations were analyzed. The solid lines correspond to the fiber conformations with NRL = 167 bp (L *≈* 10*n*), and the broken lines are for the fibers with NRL = 172 bp (L *≈* 10*n*+5). Cumulative probability P(X) represents the fraction of conformations in which the Cys-Cys distance (measured as the distance between two C*a* atoms) does not exceed a given value X. The results for NRL = 162 and 177 bp are similar to the presented distributions for NRL = 172 and 167 bp, respectively (data not shown).

Now, consider the distances between the H4-21 residues (Figure 4B). Here, the NRL = 167 bp fiber is the clear leader, with more than 30% of MC conformations satisfying the distance threshold of 30 Å (compared to ~10% for NRL = 167 bp). Again, this tendency is consistent with the experiment (Figure 2B). Overall, the predicted probability of formation of the disulfide bonds increases in the order:

H2B-H2B (167 bp) << H2B-H2B (172 bp)) *»* H4-H4 (172 bp < H4-H4 (167 bp), which is in a close agreement with the measurements by Ekundayo *et al*. [52]. Notably, for the other members of the {10n} and {10n+5} families, the relative order of predicted probabilities is the same (data not shown).

The representative fiber conformations with short Cys-Cys distances, that can serve as transient intermediates for formation of the disulfide bonds, are shown in Figure 5. The 167-bp fiber conformation is favorable for the H4-H4 crosslink, and the two adjacent nucleosomes (i) and (i+2) are in a spatial configuration predicted in Figure 3 (with the “top” nucleosome shifted to the right). On the other hand, the 172-bp conformation is suitable for the H2B-H2B crosslink (with the “top” nucleosome shifted to the left). Note that the 172-bp fiber is characterized by nucleosome stacking that is very close to that observed in the crystal ladder-like structure of the ‘601’ nucleosome in complex with the globular domain of linker histone H5 [55].

**Figure 5.**
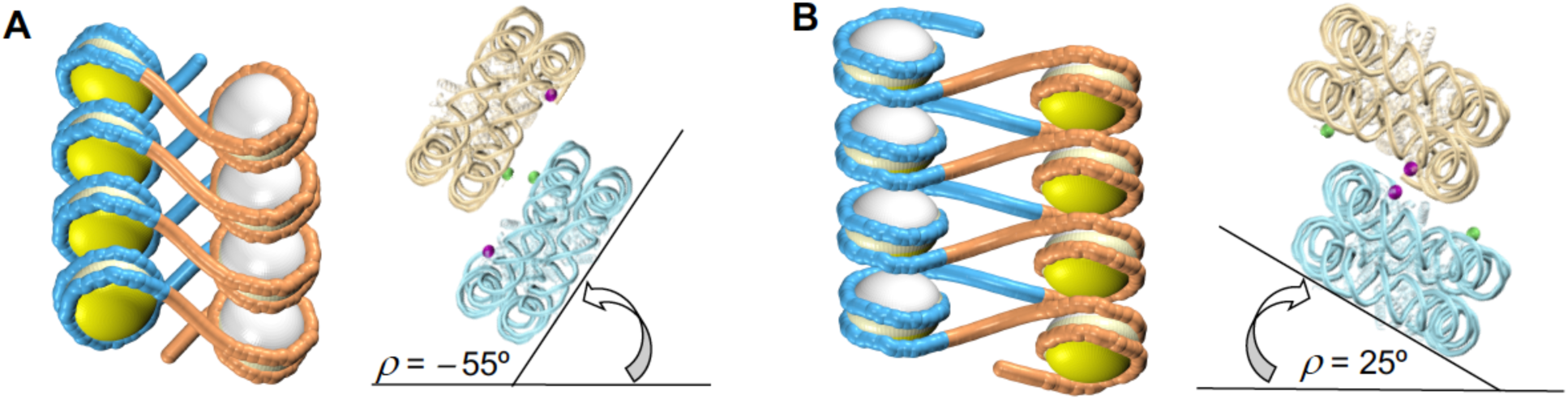
Fiber conformations with short Cys-Cys distances ~10 Å, for NRL = 167 bp **(A)** and 172 bp **(B)**. The inclination angle *r* = *-*55º for NRL=167 bp; *r* = 25º for NRL=172 bp. The fiber radius is 69 Å (A) and 85 Å (B). Note a remarkable similarity between the 172-bp conformation (B) and the fiber model based on the observed nucleosome packing in the crystal, with NRL = 171 bp (see Figure 5C in [55]).

Thus, we see that our MC simulations provide a plausible structural interpretation of the strong dependence of histone crosslinks on the nucleosome spacing, observed by Ekundayo *et al.* [52]. This remarkable consistency between our theory and experiment can be considered as yet another evidence for existence of the novel {10n+5} fiber topoisomers in solution.

## Radioprobing DNA folding *in situ*

Recently, Risca *et al*. [53] used ionizing Radiation-Induced spatially Correlated Cleavage of DNA with sequencing (RICC-seq) to identify the DNA-DNA contacts that are spatially proximal *in situ*, in human cells. The two experimentally observed dominant fragment sizes, 280-290 and 360-370 nt, are consistent with the formation of a two-start chromatin fiber similar to that resolved by Cryo-EM [8], see Figure 6A. These results reproduce the pioneering study by Rydberg *et al.* [56], who were the first to assign the peaks in RICC cleavage to the zig-zag folding of DNA in the 30-nm fiber. In addition, Risca *et al.* [53] employed the new-generation sequencing technique providing genome-wide information on the relative intensities of the RICC-seq peaks, which reflect spatial organization of chromatin fibers. Remarkably, this information can be linked to the epigenetic maps of the active and repressed states of chromatin in different parts of human genome (see below).

**Figure 6.**
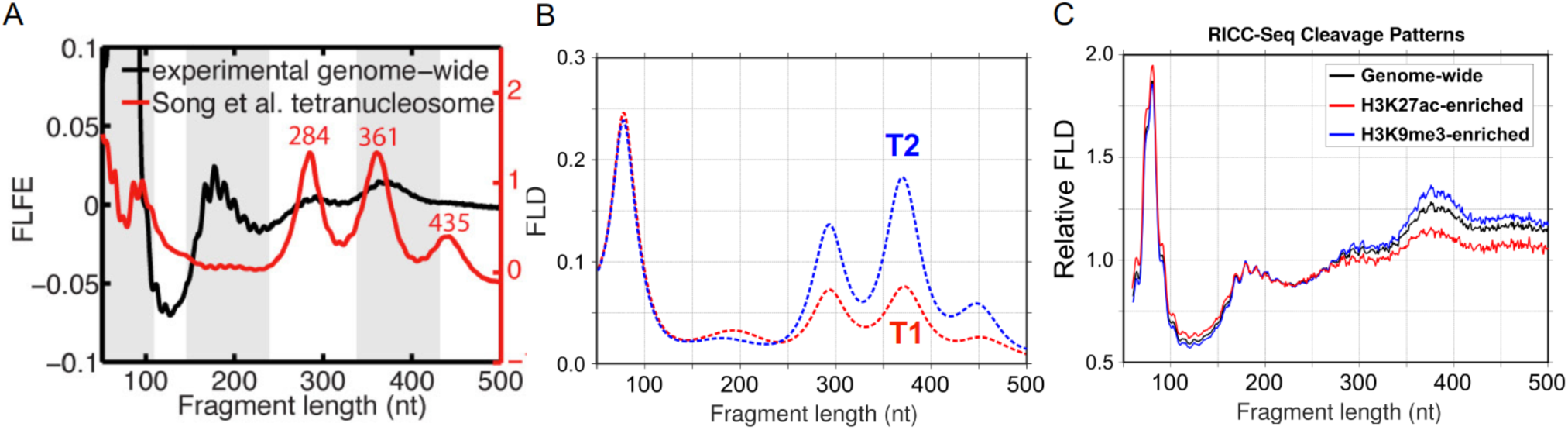
Comparison of the experimental RICC-seq data [53] with theoretical predictions. **(A)** The experimental genome-wide Fragment Length Fold-Enrichment (FLFE) profile [53] is shown by black curve. The red curve is for the Fragment Length Distribution (FLD) calculated for the Cryo-EM tetranucleosome structure with NRL = 187 bp [8]. Note that the equal heights of the 284-nt and the 361-nt peaks are not consistent with the experimental data. (Figure S10-g from Risca *et al.* [53] is reproduced with permission.) **(B)** Our MC simulated FLD profiles for the topoisomers T1 (NRL = 182 bp, in red) and T2 (NRL = 187 bp, in blue). Compare with Figure 7. **(C)** The FLD profiles calculated for the transcriptionally active (H3K27ac, red curve) and repressed (H3K9me3, blue curve) regions in human genome, based on the RICC-seq data by Risca *et al.* [53]. The genome-wide distribution is shown by black curve.

To interpret the RICC cleavage data, Risca *et al.* [53] used molecular models of the two-start and multi-start fibers generated with the software designed by Koslover *et al.* [57]. The two-start models all belong to the T2 family, containing topoisomers most favorable for the {10n} linkers [18]. However, the DNA linkers observed *in vivo* mostly belong to the {10n+5} series. In this case, the T1 topoisomer is preferable [18,34]. Therefore, we investigated how the theoretical interpretation of the RICC results would change if the two distinct topoisomers, T1 and T2, are considered.

The Fragment Length Distributions (FLDs) calculated for the T1 and T2 topoisomers (Figures 7A, B) differ in intensity of the two main peaks, at 290 and 370 nt. The FLD profile for the T2 form with NRL=187 bp is similar to that calculated by Risca *et al*. [53] for the Cryo-EM tetranucleosome structure [8], also with NRL=187 bp (Figure 6A), which is not surprising because the two conformations are topologically equivalent [34]. For the T1 topoisomer, however, the 290-nt peak is significantly lower than the 370-nt peak (Figure 7B, NRL=182 bp). This is explained by different folding of DNA in the T1 and T2 fibers (Figure 8).

**Figure 7.**
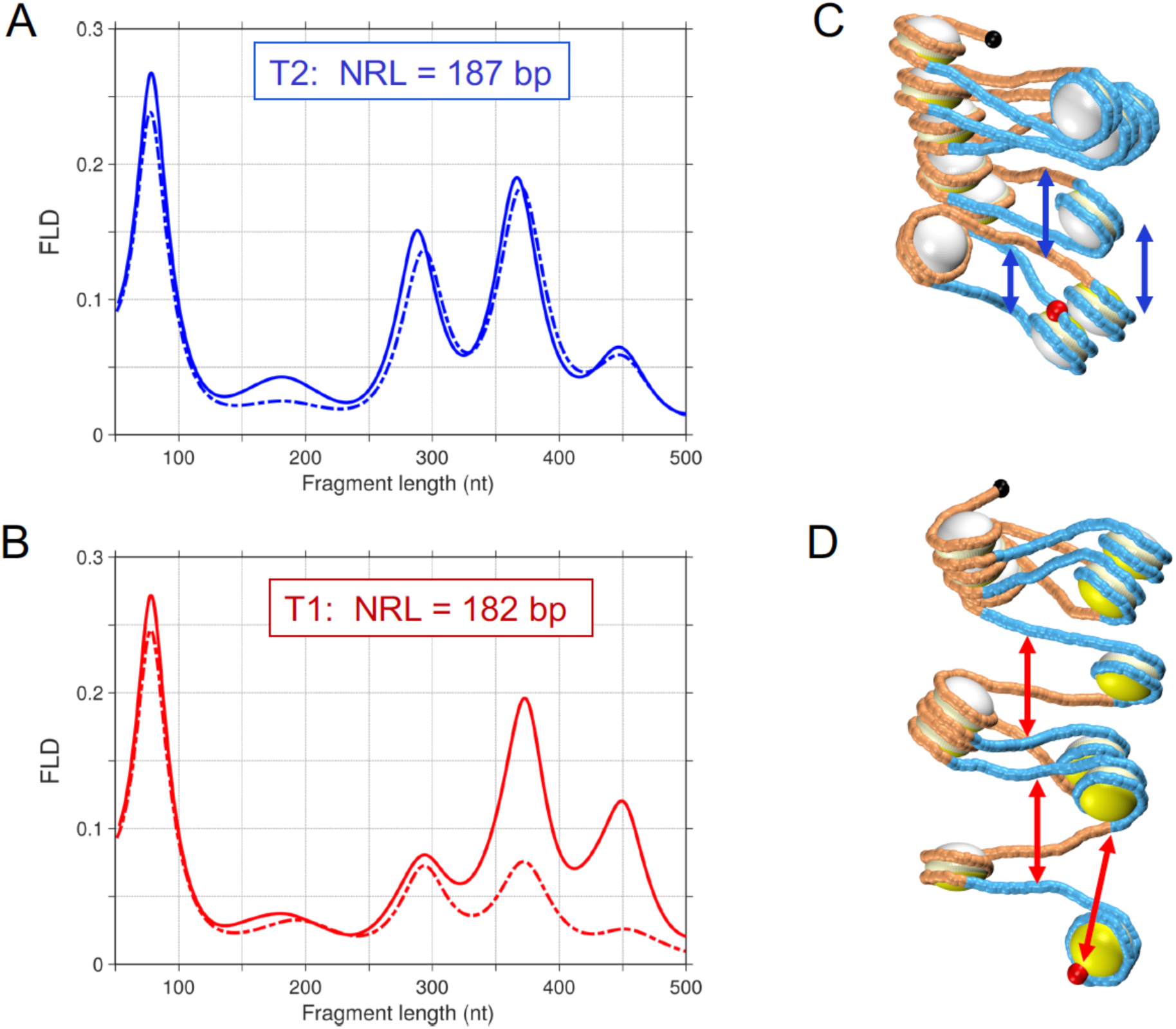
Influence of thermal fluctuations on theoretical FLD profiles. **(A, B)** The FLD curves calculated for the two-start chromatin fibers with NRL = 187 bp (A) and NRL = 182 bp (B). The data for energetically optimal structures are shown in solid lines, and for the MC ensembles, in dashed lines. The structural origin of the 180-, 290-, 370- and 450-nt peaks is described in Figure 8. The strong signal at 80 nt is due to the spatially close DNA superhelical gyres inside individual nucleosomes. To calculate FLD(*n*) for a given fiber conformation, for each fragment of length “*n*” base pairs, the correlated frequency of DNA breaks at the ends of the fragment was calculated, and then all such frequencies were summed up. The frequency of DNA breaks as a function of 3D distance between the fragment ends decreases exponentially, with the exponential drop constant *l* = 4.0 nm [53]. **(C, D)** Typical configurations of 12-nucleosome arrays with NRL = 187 bp (C) and NRL = 182 bp (D) obtained in the course of MC simulations. The arrows indicate increased distances between the DNA points separated by two nucleosome repeats (~370 nt); compare with the optimal conformations in Figure 8. The increase in distances is caused by thermal fluctuations during MC procedure [44]; it is more pronounced for NRL = 182 bp (D).

**Figure 8.**
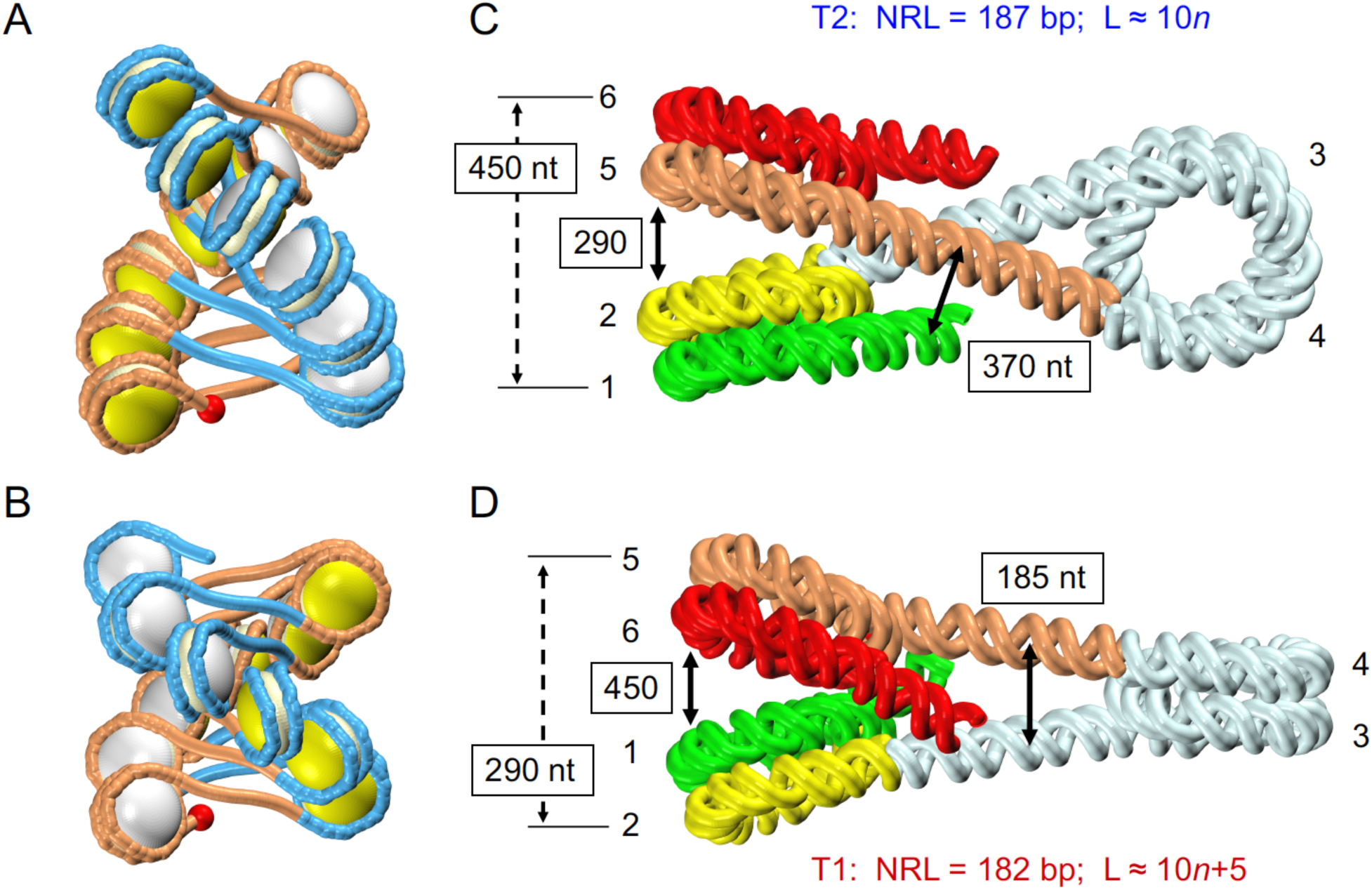
Distinct DNA folding in the T1 and T2 topoisomers leads to different FLD profiles. **(A, B)** The entry points in 12-nucleosome arrays with NRL = 187 bp (A) and NRL = 182 bp (B) are shown as red spheres. Compare with Figure 1. **(C, D)** The entry and the exit halves of nucleosomes (gyres) in the left stack are colored differently to emphasize different spatial organization of DNA in the two cases. In case of NRL = 187 bp (C), the yellow (#2) and orange (#5) gyres separated by 290 nt are spatially close. Whereas in case of NRL = 182 bp (D), the same gyres #2 and #5 are spatially distant. This explains why the 290-nt FLD peak is much stronger for NRL = 187 bp (Figures 7A, B). The situation with the gyres #1 and #6 separated by 450 nt is the opposite – the FLD peak at 450-nt is higher for NRL = 182 bp than for 187 bp (Figures 7A, B). This is also consistent with the spatial closeness of the green (#1) and red (#6) gyres in the T1 fiber with NRL = 182 bp.

In the T2 topoisomer with NRL = 187 bp (Figures 8A, C), the stacked nucleosomes are arranged in such a way that the exit of the first nucleosome (gyre #2) is close to the entry of the adjacent nucleosome (gyre #5). The two gyres are separated by 290 nt on average, therefore, the 290-nt peak is relatively strong for NRL = 187 bp (Figure 7A). By contrast, in the T1 topoisomer with NRL = 182 bp (Figures 8B, D), the gyres #2 and #5 are separated in space, therefore, the FLD peak at 290 nt is much weaker in this case (which is apparently consistent with the experimental FLD curve presented in Figure 6A). Accordingly, different intensities of the 450-nt peaks calculated for the T1 and T2 topoisomers (Figures 7A, B) can be explained by distinct spatial arrangement of the gyres #1 and #6 in the two fibers (Figures 8C, D). The other strong peak in the FLD profiles, at 370 nt (Figures 7A, B) reflects close proximity of the linkers (Figure 8C). Note that the structural assignment of all these peaks that is consistent with the two-start organization of 30-nm fiber, was initially made by Rydberg *et al*. [56].

The only remaining area where the predicted FLD profiles differ substantially from the experimental one, is the interval centered at ~180 nt (Figure 6A), corresponding to the distance between the centers of the adjacent linkers (Figure 8D). In the absence of linker histones, these DNA locations are free from direct contact with proteins, and thus, are expected to be more susceptible to ionizing radiation [58]. Our current model does not take this effect into account, but more sophisticated models are likely to overcome this discrepancy in the near future.

### Monte Carlo (MC) simulations

In addition to the “global” alteration of the fiber configuration induced by topological T2 – T1 transition, the “local” conformational flexibility has to be taken into account as well. To this aim, we performed Monte Carlo (MC) simulations of the nucleosome arrays with NRL = 182 and 187 bp. (The methodological details are given elsewhere [44].) As expected, intensities of the FLD peaks diminished compared to the energetically optimal fibers (Figures 7A, B), because the corresponding DNA-DNA distances have increased (on average) due to thermal fluctuations. This is illustrated in Figures 7C, D, where the arrows indicate the increased distances between the DNA points separated by ~370 nt (or two nucleosome repeats).

The MC-induced blurring effect proved to be the strongest for the T1 topoisomer with NRL = 182 bp (the 370-nt peak). Note that such a pronounced flexibility of the T1 fiber is in agreement with sedimentation velocity measurements [17] and our computations [18]. We also ran MC simulations for NRL = 181, 183, 186 and 188 bp. The FLD profiles deviated from the profiles for NRL = 182 and 187 bp (Figures 7A, B) by no more than 0.02 units.

Overall, the decreased FLD profile calculated for the T1 topoisomer (with NRL = 181-183 bp, L ≈ 10*n*+5) is in a qualitative agreement with the experimental data (Figures 6A, B), which may be related to the frequent occurrence of the {10*n*+5} linkers *in vivo* [36-40].

### RICC-seq data and transcription

Next, we analyzed the RICC-seq data obtained for genomic domains characterized by various levels of transcription. The active and the repressed chromatin regions are usually enriched with the H3K27ac and H3K9me3 epigenetic marks, respectively (Figure 9). The relative FLD profiles calculated for these regions based on the results of Risca *et al.* [53], are presented in Figure 6C, along with the genome-wide FLD. (See Figure 10 for details.)

**Figure 9.**
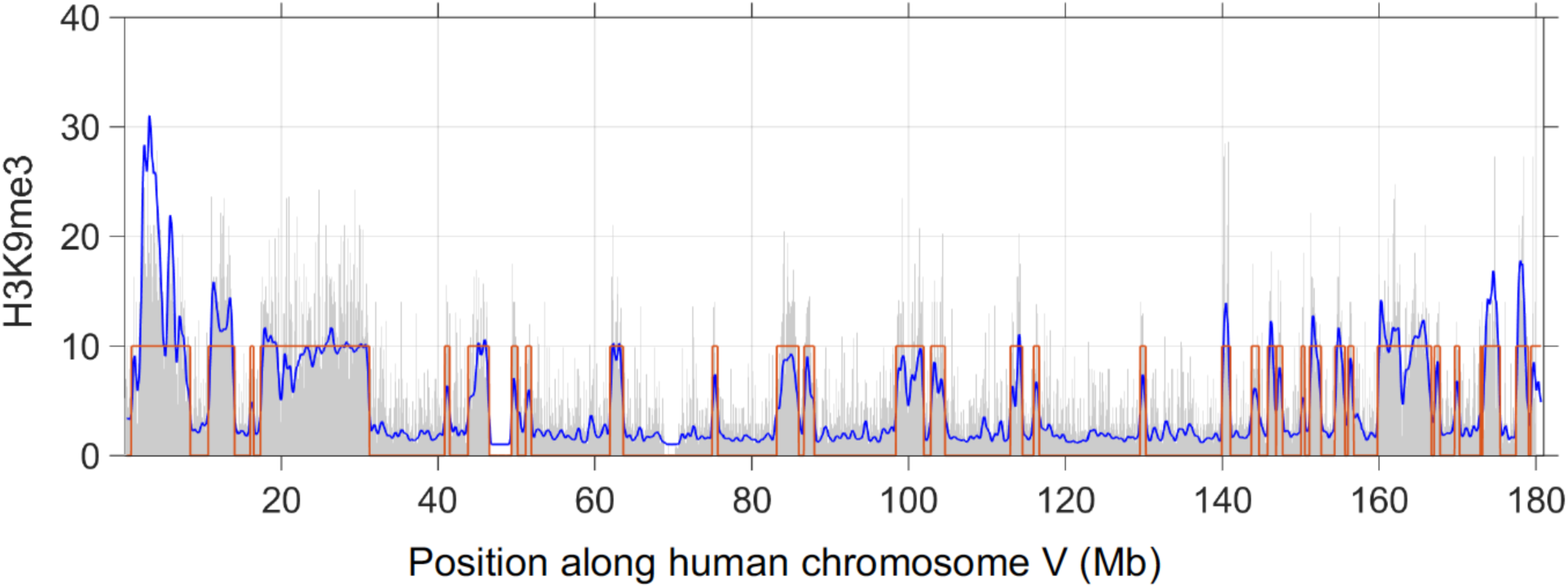
Selection of the repressed chromatin regions based on H3K9me3 ChIP-seq data [111*]. Original H3K9me3 signal shown by gray bars, is averaged using the sliding 0.5-Mb window with 50-kb increments (see the blue curve). If the average signal is higher than the threshold, the region is marked by a red rectangle. To select the active chromatin regions, the same procedure was repeated for the H3K27ac ChIP-seq data [59]. The threshold value is chosen so that ~30% of genome belongs to the regions enriched by a particular epigenetic marker. The threshold equals 4 for H3K9me3 and 3 for H3K27ac.

**Figure 10.**
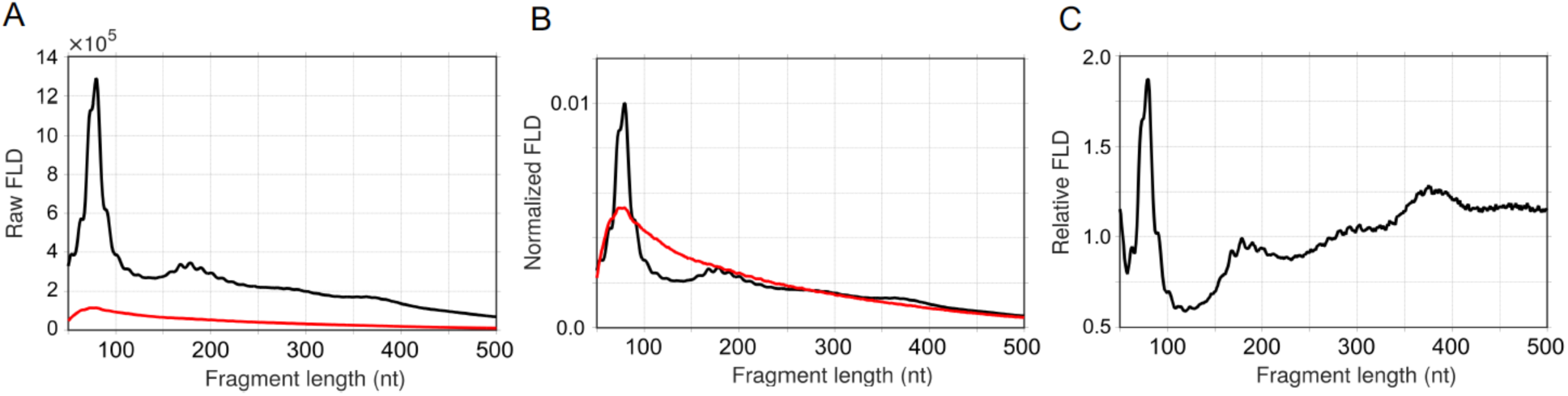
Calculations of the RICC-seq fragment length enrichment. **(A)**Genome-wide raw fragment length distributions (Raw FLDs) obtained from live irradiated cells, 300 Gy in black and 0 Gy in red [53]. The latter is used as the control background. **(B)**Fragment length distributions normalized by the total number of fragments so that the area under each curve equals 1. **(C)**Relative FLD shows enrichment in the number of DNA fragments (with a given length) in the irradiated cells compared to the control set. (The values shown in black (B) are divided by the values shown in red (B).)

These curves are clearly different, with the H3K9me3-enriched profile having the highest amplitude at ~370 nt, and the H3K27ac profile having the lowest amplitude. Naturally, the genome-wide distribution has an intermediate profile. The FLD profile generated for another epigenetic mark associated with active transcription, H3K36me3, is practically indistinguishable from the H3K27ac profile (data not shown). In a sense, the H3K9me3- and H3K27ac-enriched profiles are related in the same way as the theoretical MC profiles for the T2 and T1 topoisomers (Figure 6B). Indeed, in the T1 and H3K27ac-enriched profiles the 370-nt peak is significantly weakened (compared to the T2 and H3K9me3 profiles), such that it becomes nearly equal to the 290-nt peak.

These observations imply that transcriptionally active and repressed genomic domains are characterized by different fractions of the T1 and the T2 topoisomers, the former clearly dominating in the active domains enriched with H3K27ac mark. This is entirely consistent with our hypothesis that the novel T1 topoisomer (with L ≈ 10*n*+5) is associated with the high level of transcription [34].

In summary, relative intensities of the FLD peaks observed by RICC-seq provide important information on spatial organization of chromatin fibers in various genomic regions; in particular, they can be used to distinguish between the topological organization of the active and repressed chromatin.

## Conclusion

We have presented several lines of evidence for a topological polymorphism of chromatin fibers. In addition to the well-known T2 topoisomer, we predicted [18,34] and later observed [19] a novel T1 family of forms. The two families differ by conformational dynamics and the level of DNA supercoiling (or linking number). Importantly, the T1 and T2 topoisomers are energetically favorable for different linker lengths (L = 10*n*+5 and 10*n*, respectively). In other words, the nucleosome spacing defines topological organization and dynamics of the chromatin fiber.

The novel T1 topoisomer is characterized by an increased plasticity (which is the consequence of a weak stacking of nucleosomes). As a result, it makes chromatin more accessible to DNA binding factors and RNA transcription machinery. In addition, the T1 topoisomer has a decreased level of DNA supercoiling [19], which is usually associated with active transcription [45].

Therefore, we suggested [18,34] that the {10*n*+5} DNA linkers produce flexible and transcriptionally competent chromatin structures, while the {10*n*} linkers may be important for formation of stably folded chromatin fibers with a high level of DNA supercoiling typical of heterochromatin. This hypothesis was shown to be consistent with available data for a few species (yeast, fly, mouse and human). It remains to be seen whether the observed correlations reflect a more general tendency of chromosomal domains containing active or repressed genes (*i.e*., domains associated with different epigenetic marks) to retain topologically distinct higher-order structures.

In the last two sections we analyzed recent studies of chromatin fibers – on the nucleosome crosslinking *in vitro* [52] and on radioprobing DNA folding in human cells [53]. In both cases, we show that the novel T1 topoisomer has to be taken into account to interpret experimental data. This is yet another evidence for occurrence of alternative fiber topoisomer T1.

Our main result, the NRL-dependent topological polymorphism of chromatin fibers, can be interesting from the point of view of genome regulation and maintenance. Potentially, our findings may reveal new mechanisms for encoding structural information in the form of alternative topological states of nucleosome arrays (*i.e*., topological switch between the T1 and T2 topoisomers).

## Acknowledgements

We are grateful to Yawen Bai, William Greenleaf, Sergei Grigoryev, Viviana Risca, Igor Panyutin and Vladimir Teif for valuable discussions.

